# Click Chemistry Enables Rapid Development of Potent sEH PROTACs Using a Direct-to-Biology Approach

**DOI:** 10.1101/2025.05.19.654949

**Authors:** Julia Schoenfeld, Nick Liebisch, Steffen Brunst, Lilia Weizel, Stefan Knapp, Aimo Kannt, Ewgenij Proschak, Kerstin Hiesinger

**Affiliations:** Institute of Pharmaceutical Chemistry, Goethe University, 60438 Frankfurt am Main, Germany; Fraunhofer Institute for Translational Medicine and Pharmacology ITMP, 60596 Frankfurt am Main, Germany; Structural Genomics Consortium (SGC), Buchmann Institute for Life Sciences, 60438 Frankfurt/Main, Germany; Institute for Clinical Pharmacology, Goethe University, 60596 Frankfurt am Main, Germany

## Abstract

The Direct-to-Biology (D2B) approach enables biological screening of crude reaction mixtures, eliminating the need for purification steps and thereby accelerating drug discovery. In this study, we developed a miniaturized D2B platform for the rapid synthesis of PROTAC degraders of soluble epoxide hydrolase (sEH). We used the copper-catalyzed azide-alkyne cycloaddition and optimized the conditions for 384-well PCR plate applications with 10 µL reaction volumes on a 300 nmol scale. This approach enabled the D2B synthesis of 92 crude PROTACs from azide-functionalized CRBN-ligands and alkyne-linked sEH inhibitors. Biological screening using a HiBiT lytic degradation assay identified two hits which were resynthesized and exhibited subnanomolar DC_50_ values and degradation efficacy (D_max_). Thus, we established a scalable, cost-effective and time-saving D2B platform for the discovery of PROTACs in very small quantities. This methodology is particularly suitable for early-stage screening and hit validation assessing the degradability of a target.

## Introduction

Acceleration of the drug discovery process while ensuring reliability remains a central challenge in medicinal chemistry. One increasingly adopted solution, particularly within pharmaceutical industry,^1–3^ is the direct-to-biology (D2B) approach. This strategy involves evaluating crude reaction mixtures directly in biological assays, eliminating the need for purification steps (Figure 1). This technique dramatically reduces time and resource consumption by enabling fast hit identification, bypassing labor-intensive purification steps, and streamlining the screening workflow. The D2B approach is particularly well suited for the synthesis of proteolysis targeting chimera (PROTAC). PROTACs are heterobifunctional modalities that bind to a protein of interest and a E3 ligase to induce degradation of the protein of interest. In PROTAC discovery, hit identification requires a high synthetic effort with limited opportunities for rational design due to the complex nature of PROTAC development.^4^ Fast biological testing of the synthesized PROTACs is enabled by the HiBiT technology which allows for high throughput screening for degradation^5^, avoiding time consuming techniques as Western Blot or ELISA.

**Figure 1.**
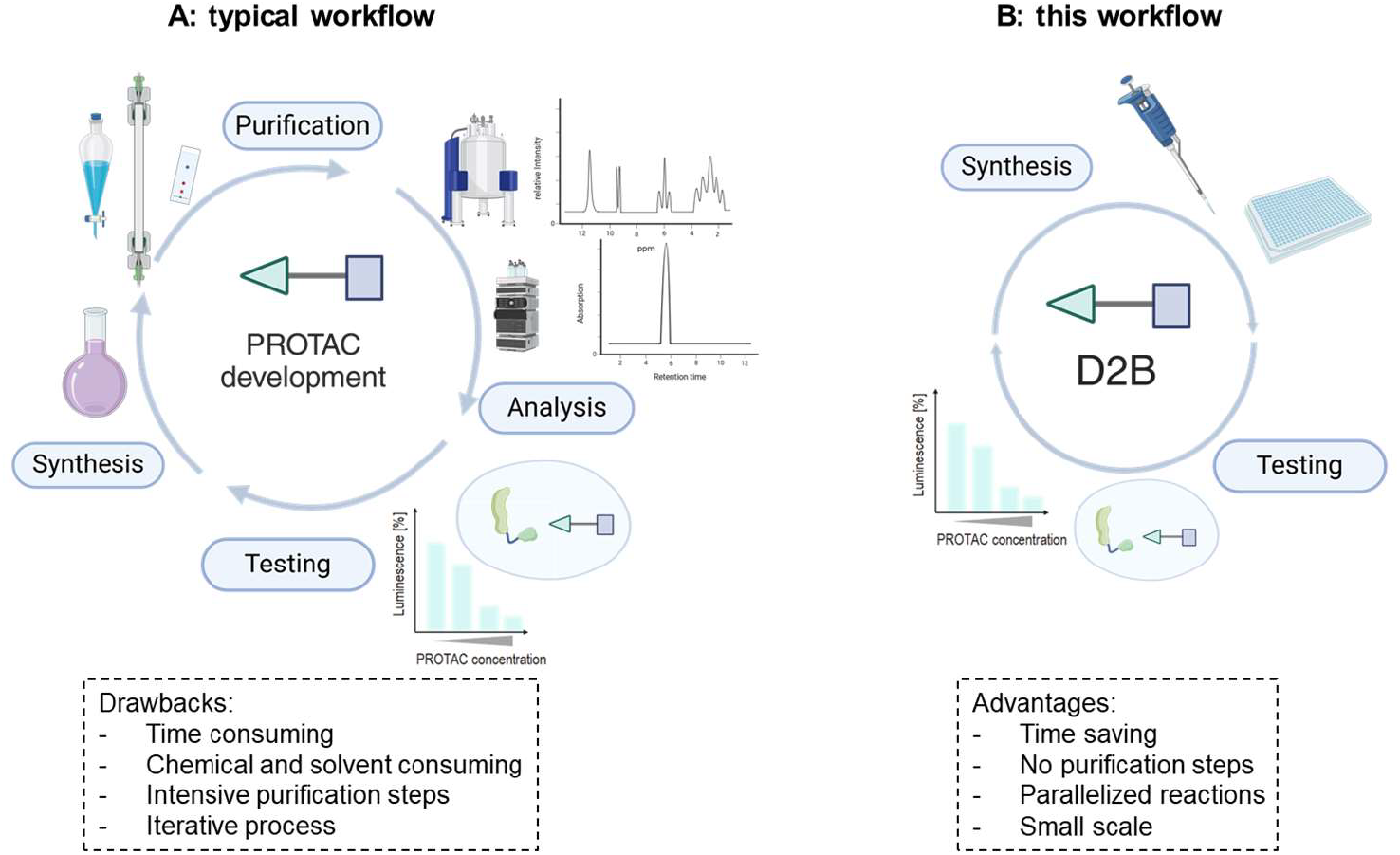
Schematic representation of two PROTAC development workflows. A: Cycle of a typical PROTAC development campaign. The steps include synthesis, purification, analysis and testing. B: Shorter workflow through D2B approach, which includes synthesis and direct testing of crude mixtures of a miniaturized PROTAC CLICK chemistry platform. The figure was prepared using BioRender.com.

The efficiency of D2B workflows is further enhanced by miniaturization of chemical reactions, which brings significant benefits such as reduced chemical waste, lower consumption of valuable starting materials and the ability to run high-throughput reactions in parallel in a plate-based format. These improvements accelerate the early stage of hit discovery while maintaining sustainability and cost efficiency. To date, most D2B platforms are based on amide coupling, as this reaction is reliable and typically results in high yields.^1–3^ Another simple and robust reaction is the copper-catalyzed azide-alkyne cycloaddition (CuAAC, CLICK chemistry), which is ideal for D2B applications as it has a high atom economy, tolerates a broad range of functional groups and requires no extremely hazardous reagents and no heating.^6^ In 2017, Wurz and coworkers demonstrated a CLICK chemistry platform for the synthesis of PROTACs (0.1 mmol scale), in which the PROTACs were evaluated for degradation towards the bromodomain extraterminal domain-4 (BRD4) after purification.^7^ Che and colleagues have recently shown that CLICK chemistry is suitable for automated synthesis in 1 mL reaction volume within a D2B approach.^8^ The present study aims at scaling down this procedure even further and optimizing the CLICK reaction for 10 µL reaction volume and nanomole scale. To evaluate the utility of our platform, we selected soluble epoxide hydrolase (sEH) as a model target due to its therapeutic relevance in the context of inflammation-related diseases^9–13^ and the availability of an in-house degradation assay utilizing HiBiT technology.^5^ sEH comprises a C-terminal epoxide hydrolase domain and a structurally distinct N-terminal lipid phosphatase domain, both of which can be concurrently eliminated via PROTAC-mediated targeted protein degradation.^14^ Notably, Peyman et al. demonstrated that an sEH-targeting PROTAC effectively reduces endoplasmic reticulum (ER) stress and inflammatory markers in hepatic cells and murine liver tissue.^15^

Here, we developed a miniaturized copper-catalyzed azide-alkyne cycloaddition in a plate-based format to accelerate hit identification in PROTACs targeting the sEH. Using this approach, we synthesized 92 crude PROTACs at the nanoscale and evaluated them with a lytic HiBiT lytic assay, leading to the identification of two hits. The resynthesized and purified PROTACs show strong degradation, demonstrating the applicability of this platform.

## Results and Discussion

In a previous study we developed sEH PROTACs using conventional combinatorial chemistry approaches and demonstrated that CRBN-based PROTACs are suitable to target sEH. Based on these data, we focused in this study on CRBN targeting ligands and obtained a CRBN-targeting azide library from Enamine containing 3 µmol of 23 azide (Scheme 1). After evaluating reported sEH PROTACs^14–17^ and crystal structures of potent sEH inhibitors, we designed four sEH inhibitors harboring a terminal alkyne oriented towards the solvent region to allow coupling with the purchased CRBN-targeting azides using CLICK chemistry.

**Scheme 1.**
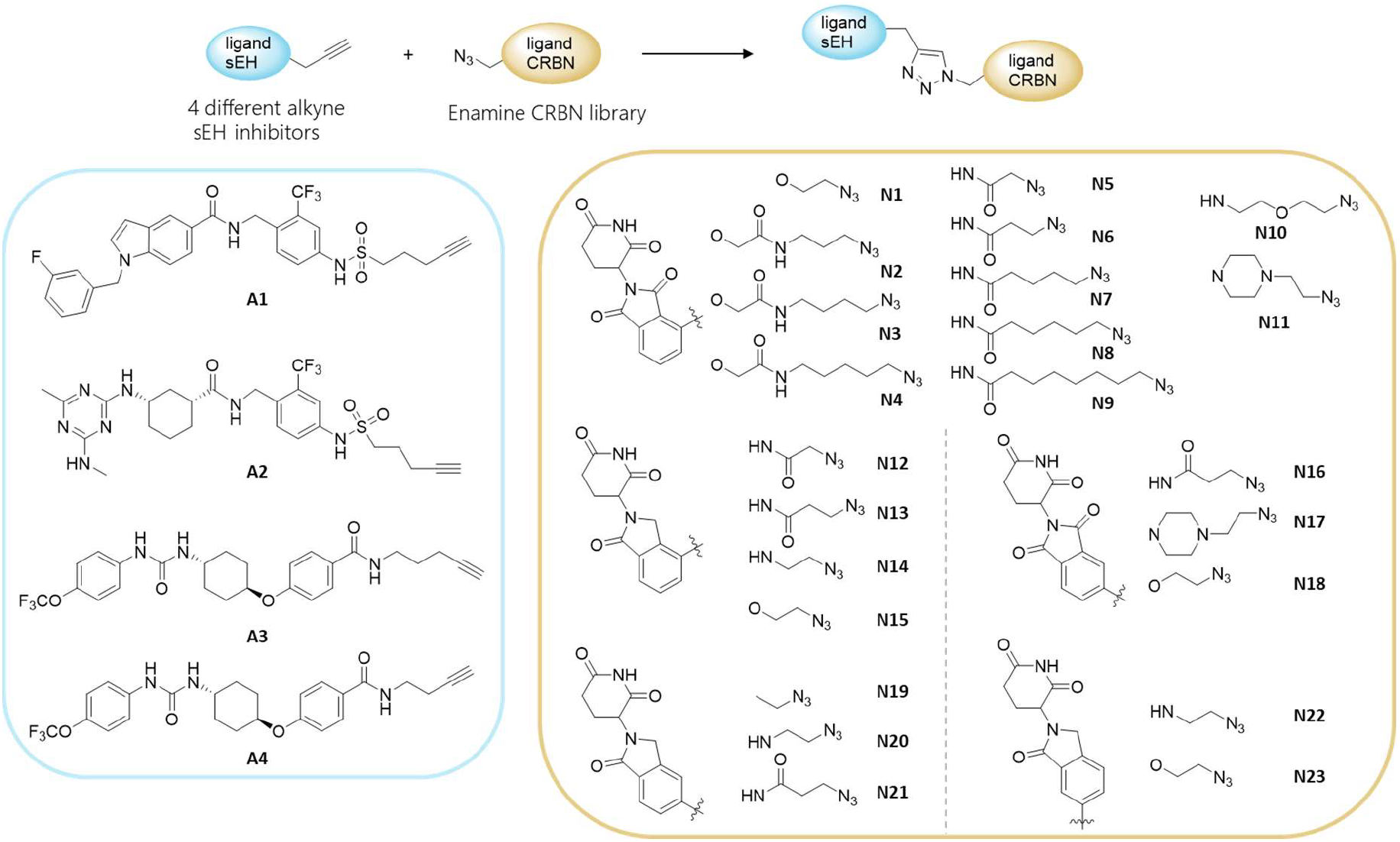
Detailed presentation of the structures of the library. Left: Chemical structures of alkyne-functionalized sEH ligands A1-A4. Right: Chemical structure of azide-functionalized CRBN ligands N1-N23.

For this study, we decided to target the C-terminal epoxide hydrolase domain (sEH-H), as sEH-H inhibitors are extensively studied and synthetically easily accessible. sEH-H exhibits an L-shaped binding pocket with two branches (short and long), both of which can be addressed for PROTAC design using an alkyne linker.^14^ We chose the sEH-H inhibitors t-TUCB and FL217, as these ligands were already successfully used for PROTAC design as well as GSK2256294 which is a highly potent sEH-H ligand with excellent pharmacokinetic properties.^18–20^ For the exit of the short branch, the FL217-based ligand **A1** and the GSK2256294-based ligand **A2** were selected, while the t-TUCB-based ligands **A3** and **A4** were designed for the long branch (Scheme 1). For the optimization studies, we chose the reaction of sEH ligand **A1** with Pomalidomide-PEG5-azide (**N24**) or Thalidomide-*O*-amido-PEG4 (**N25**) to PROTAC **P1** or **P2**, respectively, which has been successful in our previous study. We explored that using CuI in dichloromethane did not lead to full reaction conversion (Table 1, entry 1). Switching to CuSO_4_·5H_2_O and sodium ascorbate as reducing agent improved the conversion (entry 2,3). Increasing the amount of the catalytic system and exchanging the solvent to a DMF and water mix gave the desired product with high conversion rates and acceptable isolated yield (entry 5). Initially, the reaction was carried out in an Eppendorf tube in 130 µL of DMF/water (4:1). To avoid using a hazardous solvent such as DMF, we decided to use DMSO. DMSO has the advantages of a high boiling point to avoid concentration fluctuations in the planned plate-based reactions and of being biocompatible. With these optimized reaction conditions in hand, we aimed to minimize the scale of the reaction. First optimization experiments for down-scaling the reaction were performed in a 96-well plate with a reaction volume of 50 µL and a concentration of 60 mM ligand **A1**, where only a moderate conversion was observed by HPLC (Table 1, entry 7). Using a 384-well plate with the same volume restored a high conversion rate, probably due to a more favorable solvent surface area (entry 8). Next, we reduced the concentration of the reactants systematically and found that the CLICK reaction is highly concentration-dependent, as previously reported by Hasimoto et al.^21^ When using the reactants in a lower concentration, the conversion decreased (Figure 2A). For this reason, we kept a concentration of 60 or 30 mM (relative to starting material) and reduced the reaction volume down to 5 µL. We used a 384-well plate to achieve a smaller solvent surface area and better mixing of the reactants. Conversion rate was higher in 384-well plates with a v-shaped bottom compared to flat bottom plates (Figures 2B and 2C). With v-shaped 384-well PCR plates, we were able to recover the conversion also at lower reaction volumes. At 5 µL and 10 µL reaction volumes with 150 nmol and 300 nmol alkyne/azide, respectively, high conversion rates (98 and 97 %, respectively) were achieved (Figure 2C). To test the robustness of the system, we ran the reaction at least three times for each volume and found pipetting small volumes accurately by hand to be challenging (Figure 2D).

**Table 1:** Optimization study of the CuAAC in small volumes.

| Entry | n | Eq. Azide | Cu(I) source | Additive | μL | solvent | time | conversion alkyne:product | yield |
| --- | --- | --- | --- | --- | --- | --- | --- | --- | --- |
| 1 | 2 | 1.05 | 20 mol% CuI | 40 mol% DIPEA, 40 mol% HOAc | 1000 | CH<sub>2</sub>Cl<sub>2</sub> | 2 h | 58:42 | 20% |
| 2 | 2 | 1.05 | 10 mol% CuSO<sub>4</sub>·5 H<sub>2</sub>O | 10 mol% Ascorbate, 10 mol% PhCO<sub>2</sub>H | 1000 | <sup>t</sup>BuOH:H<sub>2</sub>O 1:2 | 2 h | 50:50 | 25% |
| 3 | 2 | 1.20 | 10 mol% CuSO<sub>4</sub>·5 H<sub>2</sub>O | 40 mol% ascorbate | 1600 | <sup>t</sup>BuOH:H<sub>2</sub>O 1:1 | 24 h | 10:90 | 45% |
| 4 | 2 | 1.10 | 0.3 eq CuSO<sub>4</sub>·5 H<sub>2</sub>O | 0.3 eq ascorbate | 1300 | DMF:H<sub>2</sub>O 4:1 | 16 h | 2:98 | 45% |
| 5 | 4 | 1.10 | 0.3 eq CuSO<sub>4</sub>·5 H<sub>2</sub>O | 0.3 eq ascorbate | 1300 | DMF:H<sub>2</sub>O 4:1 | 16 h | 2:98 | 42% |
| 6 | 4 | 1.00 | 0.3 eq CuSO<sub>4</sub>·5 H<sub>2</sub>O | 0.3 eq ascorbate | 50 | DMF:H<sub>2</sub>O 4:1 | 16 h | 2:98 | 49% |
| 7<sup>a</sup> | 5 | 1.00 | 0.3 eq CuSO<sub>4</sub>·5 H<sub>2</sub>O | 0.3 eq ascorbate | 50 | DMSO:H<sub>2</sub>O 4:1 | 16 h | 17:82 | - |
| 8<sup>b</sup> | 5 | 1.00 | 0.3 eq CuSO<sub>4</sub>·5 H<sub>2</sub>O | 0.3 eq ascorbate | 50 | DMSO:H<sub>2</sub>O 4:1 | 16 h | 2:98 | - |

<sup>a</sup> reaction was performed in a 96-well plate <sup>b</sup> reaction was performed in a 384-well plate

**Figure 2.**
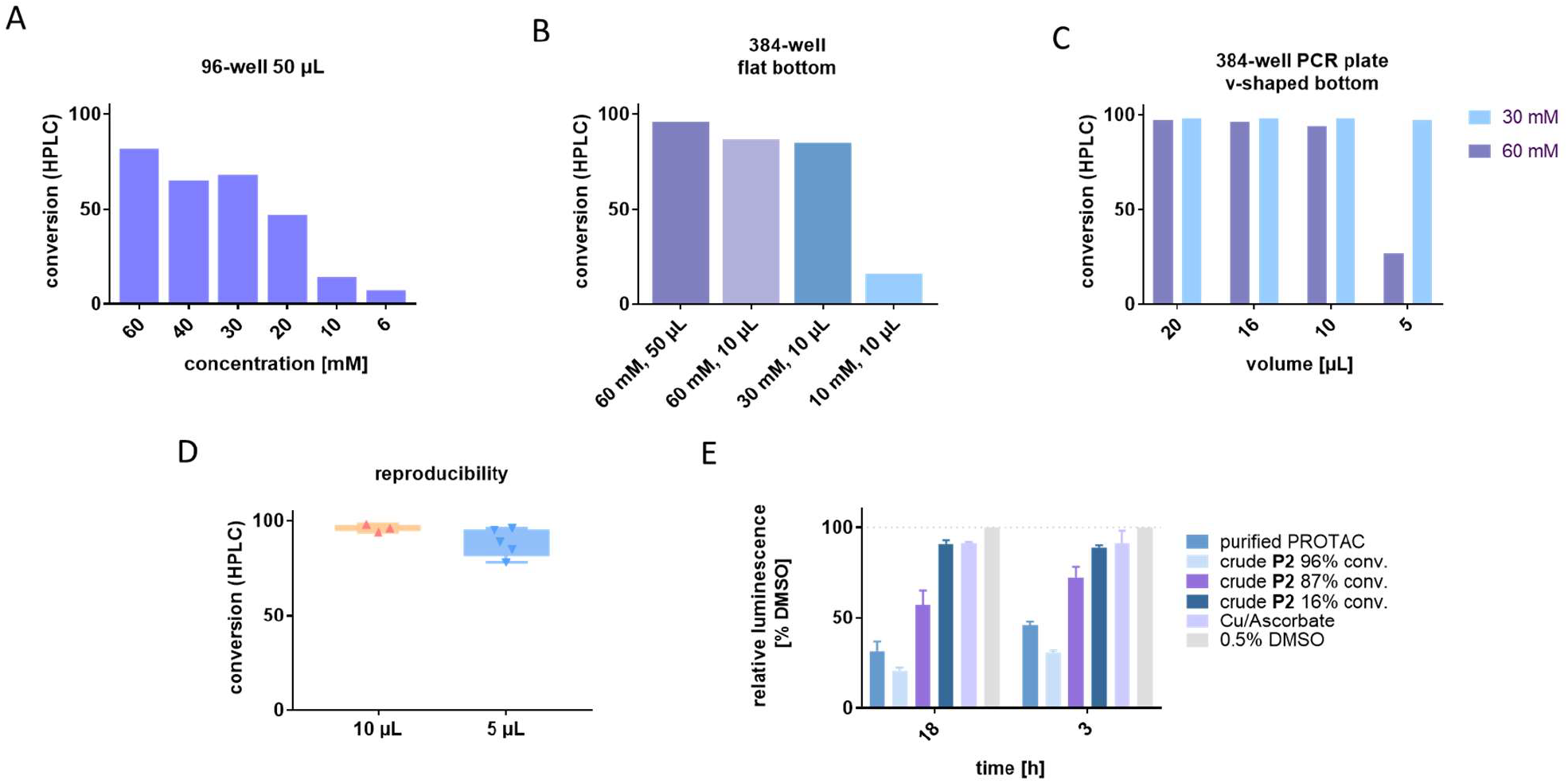
Optimization studies of the CLICK reactions. A: Concentration dependency of CuAAC reaction; conversion determined by integration of the signal of the HPLC absorption traces (254 nm). B: Conversion of CuAAC reactions with decreasing reaction volumes in a 384-well plate by integration of the signal of the HPLC absorption traces (254 nm). C: Comparison of conversion at 60 and 30 mM concentration of starting material in a 384-well plate. D: Reproducibility of the CLICK reactions in 5 µL and 10 µL reaction volume. E: Purified and crude mixtures of PROTACs were tested in a HiBiT lytic degradation assay. Cells were incubated for 18 h or 3 h with 300 nM PROTAC/crude mixtures.

In parallel, we tested the crude reaction mixtures of our model system (PROTAC **P2**) with our optimized conditions (3 h and 18 h incubation, 300 nM PROTAC/crude) in our previously reported HiBiT lytic assay system (Figure 2E).^14^ In this cell-based assay, bioluminescence is used to measure the degradation of sEH. The sEH protein is tagged with a small peptide fragment of Nanoluciferase (NanoLuc) called HiBiT. With addition of the LgBiT peptide, which marks the residual scaffold of NanoLuc, and the substrate, bioluminescence is generated. The more sEH is degraded, the less bioluminescence is detected. We found that less sEH was degraded using PROTACs synthesized with lower conversion rates. This was most likely due to a lower resulting concentration of the respective PROTAC in solution and because the unreacted alkyne may have competed with the PROTAC for binding to sEH. As copper species may occur cytotoxic, we tested the influence of the catalytic system of copper sulfate and sodium ascorbate on the HiBiT cells in the concentration present in the assay and found only very slight changes in luminescence of maximum 5% (Figure 2E). Furthermore, no cytotoxicity was observed in a cell viability test of the HiBiT cell line at the copper concentration present in the HiBiT lytic assay (Supplementary Figure 1).

With the optimized reaction conditions in hand, we performed the parallel synthesis of 92 PROTACs from the sEH ligands functionalized with an alkyne **A1-A4** and the purchased azide library (Scheme 1). We prepared DMSO stock solutions for the azide- and alkyne-functionalized ligands and water stock solutions for CuSO_4_·5H_2_O and sodium ascorbate. The respective volumes of azide, alkyne, copper(I) source and reducing agent were added to the 384-well PCR plate and the plate was sealed with aluminum foil. The plate was shaken overnight on a plate shaker, and 1 µL of each reaction mixture was used to determine the conversion. Another 1 µL of the crudes was used to prepare the stocks for biological evaluation (Figure 3A). In this setup, we observed a mean conversion of 85%, with the lowest conversion being 28%. We tested all crude samples with a 300 nM concentration in the HiBiT lytic assay, as this was an effective PROTAC concentration in our previous study, and were able to detect degradation of sEH screening the crude reaction mixtures (Figure 3 B-E). As a hit threshold, we chose a degradation of at least 50%. To investigate whether we could reproduce the identified hits, we tested the synthesized CLICK library a second time in the HiBiT lytic assay. We were able to observe the same degradation pattern and identified the same hits. Next, we ran the CLICK reaction using the same library a second time to see if we would be able to reproduce the hits. Since no hits were obtained with **A2** and **A4**, we only used **A1** and **A3** for the second run of the CLICK reaction. In this experiment, we observed a lower conversion, with a median conversion of 77% (Supplementary Figure 2). This is due to two azides not being transferred to the reaction plate, leading to only the alkynes to be detectable in the HPLC traces. Excluding these two reactions resulted in an average conversion of 82%, which was almost identical to the conversion rate observed in the first experiment. We were able to confirm the identified hits in the HiBiT lytic assay as before. We obtained two hits with only one sEH scaffold (**A3**). PROTACs **P3** and **P4**, which are combinations of sEH ligand **A3** and **N23**, a lenalidomide-derived CRBN ligand with a short alkyl chain as linker, and **N17**, a pomalidomide substructure with a piperazine moiety in the linker. To confirm our hits, we resynthesized **P3** and **P4** (Figure 4A), purified the compounds by preparative HPLC, and evaluated them in the HiBiT lytic assay. We measured the dose-response at different incubation times and determined the DC_max_ and DC_50_. **P3** exhibited a pDC_50_ of 9.17± 0.06 and a D_max_ of 91 ± 2% and **P4** a pDC_50_ of 10.3 ± 0.2 and a D_max_ of 96 ± 1%, both after 18 h incubation (Figure 4 B, C). In a spiking experiment we added the sEH ligand **A3** to the pure PROTACs (Figure 4D) and observed the same effects as in our first experiments (Figure 2E). Less degradation was observed with higher amounts of **A3** as it outcompetes the PROTAC from the binding site of sEH-H, which could potentially lead to false negative results. However, in the case of highly potent PROTAC **P4** we were still able to detect this hit, even though the conversion was only moderate.

**Figure 3.**
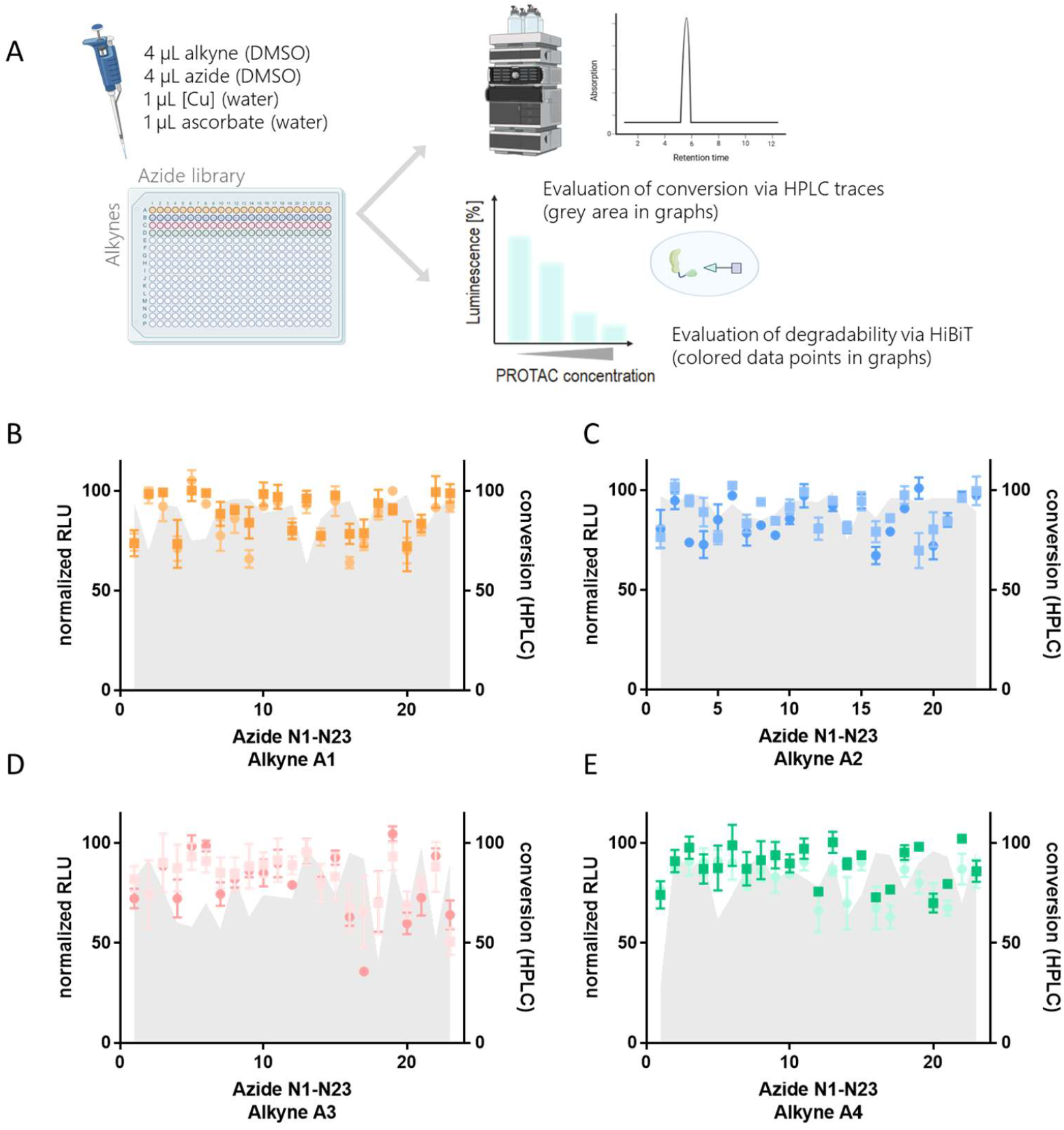
Results of the D2B approach. A: Optimized conditions for the miniaturized PROTAC CLICK platform and the post-reaction workflow. The conversion was determined by signal integration of the absorbance at 254 nm in the HPLC spectrum and in parallel sEH degradation was evaluated in a cell-based HiBiT lytic assay. B-E: Degradation activities of the crude mixtures in a HiBiT lytic assay. Cells were treated with 300 nM of the crude mixtures for 18 h (n=2 in triplicates, error bars in standard deviation (SD)); the grey area in the graph represents the conversion determined by HPLC. The figure was prepared using BioRender.com.

**Figure 4.**
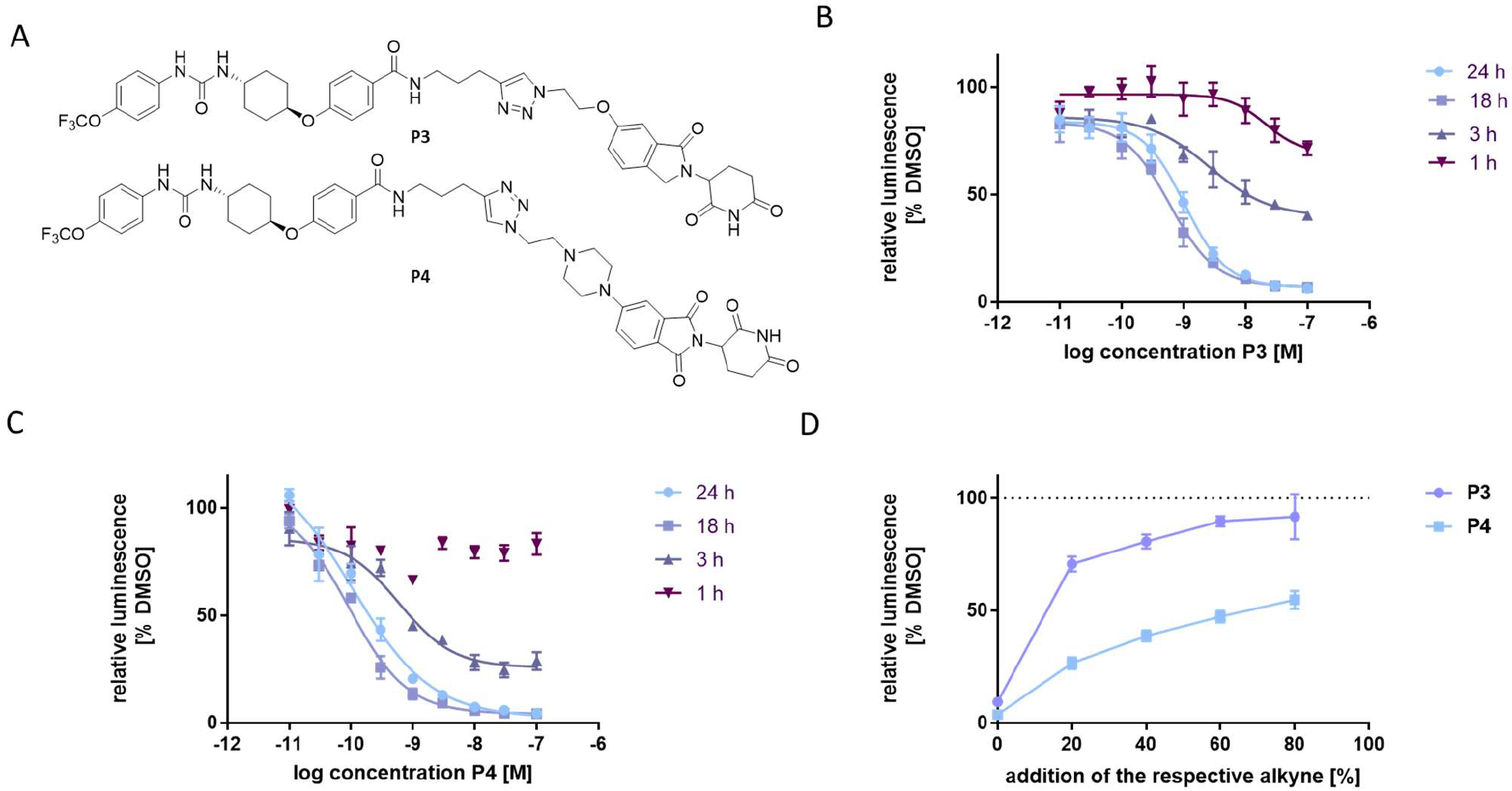
Detailed analysis of the purified hits. A: structures of PROTAC hits. B: time-dependent degradation ability of purified hit **P3** (n=2 in triplicates, error bars in SD). C: time-dependent degradation ability of purified hit **P4** (n=2 in triplicates, error bars in SD). D: Spiking experiments with addition of alkynes **N17** or **N23** to the purified PROTACs **P3** or **P4**, respectively (n=2 in triplicates, error bars in SD).

## Conclusion

In this study, we successfully developed and optimized a highly miniaturized D2B platform for the synthesis and biological evaluation of PROTACs targeting sEH, utilizing CLICK chemistry (CuAAC reaction). Our results demonstrate that the click reaction can be effectively scaled down to reaction volumes as low as 10 µL while maintaining high conversion rates, provided appropriate plate formats and reagent concentrations are used. By applying this workflow to a thaliomide-based azide library and alkyne-functionalized sEH inhibitors, we achieved the parallel synthesis of 92 unpurified potential PROTACs. Direct screening of crude reaction mixtures in a HiBiT lytic degradation assay led to two hits which were resynthesized. These PROTACs exhibited low nanomolar to subnanomolar DC_50_ values and degraded sEH with a Dmax of 90-96%, validating the biological relevance of the hits despite moderate reaction conversions in some cases. However, we also observed that residual unreacted alkyne-functionalized sEH ligand can interfere with degradation activity, underscoring the importance of reaction completeness in D2B workflows to avoid false negatives. Overall, our findings establish a scalable, cost-effective, and time-efficient platform for the discovery of PROTAC degraders in ultra-low volumes. This methodology is particularly suited to evaluate the “PROTACability” of a protein of interest for which ligands are available, for early-stage screening and hit validation as we could detect two new potent PROTACs within few weeks. However, in this study we relied on a purchased small CRBN ligand library with only moderate complexity. To increase diversity and complexity in the linker and E3 ligase ligands a more diverse library will be established and evaluated in our group in the future.

## Methods

### General Materials

Starting materials, reagents and solvents were purchased from Sigma-Aldrich (Merck), abcr, Fluka, BLDpharm, and FluoroChem and used without further purification. The azide library used for miniaturized CuAAC reactions was purchased from Enamine (CRBN Azide Kit-1, #Q1790763_EUR). Solvents were of reagent grade or dry, if specified. ALUGRAM® silica gel sheets purchased from Macherey-Nagel were used to perform thin-layer chromatography, spots were visualized by UV light (245 nm and 365 nm). Flash-Chromatography was performed on a puriFlash XS 420 device from interchim with a SPD-20A UV/VIS detector from Shimadzu. For normal phase chromatography, technical grade solvents were used as well as Silica HP 30 μm columns from interchim (size F0012, F0025 or F0040, depending on the scale of the reaction). Reversed phase chromatography was conducted using C18 HP 30 μm columns from interchim (size F0012, F0025 or F0040, depending on the scale of the reaction), eluents were ACN (HPLC grade) and milli-Q purified water. NMR spectra were recorded on either a Bruker Avance-300, Bruker Avance-400 or Bruker Avance-500 spectrometer. Spectra were calibrated on the respective non-deuterated solvent residue peak. ^1^H NMR data are reported with the chemical shift δ (ppm) relative to tetramethylsilane, integrals, multiplicity and coupling constant (Hz). For ^13^C NMR and ^19^F NMR signals chemical shifts δ (ppm) are reported. HPLC-MS analysis was performed with a flow rate 1 mL/min on a Shimadzu prominence separation device (column Luna 10 μC18(2) 100A (250 ×4.60 mm) by Phenomenex) connected with a Shimadzu SPD-20A UV/VIS UV-detector (monitoring at 254 nm and 280 nm) and a Shimadzu LCMS-2020 Single Quadrupole mass spectrometer with electrospray ionization (ESI). PROTACs **P3** and **P4** were purified using a Shimadzu prominence preparative HPLC system (column Luna 10 µC18(2) (250·21,20 nm) by Phenomenex) with UV monitoring at 254 nm and 280 nm and a flow rate 21 mL/min. For both analytical and preparative HPLC the mobile phase consisted of ACN/0.1% aqueous formic acid. Gradient for method A: 70% 0.1% aqueous formic acid for 10 min, 5 min of lowering 0.1% aqueous formic acid to 10%. Gradient for method B: 95% 0.1% aqueous formic acid for 10 min, 5 min of lowering 0.1% aqueous formic acid to 10%. Precision mass measurements were measured with ESI on a Bruker MicrOTOF-Q II mass spectrometer. Theoretical m/z and exact mass were calculated using Chemcalc.org.^22^ Purity is > 95% of all tested compounds determined through HPLC-MS analysis. sEH ligand **A1** was synthesized according to a published method.^14^ Synthesis and analytical data for PROTACs **P1** and **P2** was as described previously.^14^

## Synthesis Procedures

### Parallel synthesis of 96 sEH PROTACs

In preparation for the reactions, 75 mM DMSO stocks were prepared of every sEH ligand (alkynes) as well as every CRBN ligand (azides) from the CRBN Azide Kit-1 purchased from Enamine. After preparing the azide stocks in Eppendorf tubes, they were transferred into a 96-wel lplate, sealed with aluminium foil and stored at - 20 °C. The alkyne stocks were stored in Eppendorf tubes at -20 °C. For every reaction series, 90 mM stocks in water were prepared for CuSO_4_·5 H_2_O and sodium ascorbate, respectively. The reactions were performed in a 384-well PCR-plate (# 12677546) from Thermo Scientific. First, 23 x 4 µL of each alkyne solution (A1-A4) were transferred to the plate using an E1-ClipTip™ multichannel micropipette from Fischer Scientific (# 15663046, 2-125 µL) and the plate was centrifuged for 30 s at 500 rpm. Then, the same E1-ClipTip™ multichannel micropipette was used to transfer 4 µL of each azide solution (N1-N23) to the respective wells containing the alkynes, followed by centrifugation of the plate for 30 s at 500 rpm (final concentration of the reactants: 30 mM, 1.0 eq). In the final two steps, 1 µL of the CuSO_4_·5 H_2_O and sodium ascorbate stock solutions respectively were added to every well (final concentration of the catalyst/additive: 9 mM, 0.3 eq), using an E1-ClipTip™ multichannel micropipette from Fischer Scientific (#15623046, 0.5-12.5 µL). Afterwards, the plate was centrifuged for 1 min at 500 rpm. The plate was sealed with aluminium foil and shaken at 300 rpm for 16 h at rt on a Thermo-Shaker PHMP-4 from Grant-bio. After the reaction time, 1 µL of every well was used to prepare a 100 fold diluted solution in MeOH (HPLC-grade) for LC-MS analysis, where conversion was determined by integration of the signal of the absorption traces (254 nm). Another 1 µL was taken from every well and diluted 500 fold in DMSO for testing in the sEH HiBiT lytic assay.

### Alkyne Synthesis

*tert-*butyl-*((1S*,*3R)-*3-((4-amino-2(trifluormethyl)benzyl)carbamoyl)cyclohexyl)carbamate*: (*1R*,*3S*)-3-((*tert*-Butoxycarbonyl)amino)cyclohexane-1-carboxylic acid (150 mg, 0.78 mmol, 1.0 eq), PyBOP (447 mg, 0.86 mmol, 1.1 eq), and DIPEA (0.41 mL, 2.34 mmol, 3.0 eq) were dissolved in dry THF (15 mL) and stirred for 10min at rt Afterwards, HOBt . H_2_O (60 mg, 0.39 mmol, 0.5 eq) and 4-(aminomethyl)-3-(trifluoromethyl)aniline (200 mg, 0.78 mmol, 1.0 eq) were added to the reaction and the mixture was stirred for 16 h at rt The solution was washed with brine (2 x 15 mL) and extracted with EtOAc (3 x 20 mL). The combined organic phase was dried over MgSO_4_, filtered and concentrated in vacuo. Subsequently, the orange-yellow crude product was purified using flash chromatography (*n*-Hex:EtOAc = 100:0 → 70:30) to obtain a yellow solid in 82 % yield (266 mg, 0.64 mmol). ^1^H NMR (250 MHz, CDCl_3_): *δ* 7.27 (d, J = 8.2 Hz, 1H), 6.91 (d, J = 2.5 Hz, 1H), 6.76 (dd, J = 2.4 Hz, J = 8.2 Hz, 1H), 5.72 (t, J = 5.4 Hz, 1H), 4.45 (s, 2H), 4.43 (s, 1H), 3.85 (s, 2H), 3.48 -3.38 (m, 2H), 2.13 -2.07 (m, 2H), 1.95 -1.90 (m, 2H), 1.86 -1.76 (m, 1H), 1.82 -1.76 (m, 2H), 1.42 (s, 9H), 1.08 -1.02 (m, 1H). LC-MS (ESI) m/z: [M+H]^+^calcd. for C_20_H_29_F_3_N_3_O_3_^+^, 416.21; found, 416.15.

*tert*-butyl(*(1S*,*3R)-*3-((4-(pent-4-yn-1-ylsulfonamido)-2-(trifluoromethyl)benzyl)carbamoyl)-cyclohexyl)carbamate: *tert*-butyl-((*1S*,*3R*)-3-((4-amino-2-(trifluoromethyl)benzyl)carbamoyl)cyclohexyl)-carbamate (100 mg, 0.24 mmol, 1.0 eq) was dissolved in dry CHCl_3_ (20 mL) and sparged with argon for 15 min. Pent-4-yn-1-sulfonyl chloride (48 mg, 0.27 mmol, 1.1 eq) and pyridine (0.1 mL, 1.20 mmol, 5.0 eq) were added subsequently. The reaction mixture was stirred under reflux for 48 h in an argon atmosphere. Afterwards, the solution was washed with 1M hydrochloric acid (2 x 20 mL) and brine (20 mL). The aqueous phase was extracted with EtOAc (3x 20 mL) and the combined organic phases were dried over MgSO_4_ and filtered. After removing the solvent under reduced pressure, the crude product was purified using flash chromatography (n-Hex:EtOAc = 100:0 → 80:20). The product was obtained as a pale brown oil in 37 % yield (99 mg, 0.18 mmol). ^1^H NMR (400 MHz, CDCl_3_): *δ* 8.36 (s, 1H), 7.50 (d, J = 1.7 Hz, 1H), 7.36 -7.34 (m, 1H), 7.31 -7.29 (m, 1H), 6.32 -6.28 (m, 1H), 4.56 (dd, J = 7.1 Hz,J = 17.6 Hz, 2H), 4.46 (dd, J = 5.4 Hz, J = 15.7 Hz, 1H), 3.47 (s, 2H), 3.24 -3.20 (m, 2H), 2.34 (dt, J = 2.6 Hz, J = 6.7 Hz, 2H), 2.29 -2.23 (m, 1H), 2.14 - 2.11 (m, 1H), 2.02 (q, J = 7.2 Hz, 2H), 1.95 (t, J = 2.6 Hz, 1H), 1.90 -1.86 (m, 1H), 1.86 -1.78 (m, 2H), 1.43 (s, 9H), 1.31 -1.29 (m, 2H), 1.14 -1.04 (m, 1H). LC-MS (ESI) m/z: [M-H]^-^ calcd. for C_20_H_33_F_3_N_3_O_5_S^-^, 544.21; found, 544.05.

*(1R,3S)-*3-Amino-*N*-(4-(pent-4-in-1-ylsulfonamido)-2-(trifluormethyl)benzyl)cyclohexan-1-carboxamid: To a solution of *tert*-butyl((*1S,3R*)-3-((4-(pent-4-yn-1-ylsulfonamido)-2-(trifluoromethyl)-benzyl)carbamoyl)cyclo-hexyl)carbamate (83 mg, 0.15 mmol, 1.0 eq) in DCM (2.5 mL) was added TFA (0.3 mL, 7.61 mmol, 50.0 eq) and the reaction mixture was stirred for 16 h at rt After the reaction was completed, a pH of 8 was adjusted by the addition of saturated aqueous NaHCO_3_ solution. The mixture was extracted with EtOAc (5x 10 mL) and the combined organic phases were dried over MgSO_4_ and filtered. The solvent was removed under reduced pressure and the product was obtained as a colorless solid with a yield of 99 % (68 mg, 0.15 mmol) without further purification. ^1^H NMR (400 MHz, MeOH-d_4_): *δ* 7.56-7.55 (m, 1H), 7.47-7.42 (m, 2H), 4.91 (s, 2H), 3.24 -3.20 (m, 2H), 3.15 (tt, J = 11.7 Hz, J = 3.92 Hz, 1H), 2.45 (tt, J = 11.8 Hz, J = 3.5 Hz, 1H), 2.31 (td, J = 6.9 Hz, J = 2.3 Hz, 2H), 2.21 (t, J = 2.6 Hz, 1H), 2.12 -2.01 (m, 2H), 1.98 -1.90 (m, 2H), 1.94 -1.86 (m, 2H), 1.47 -1.27 (m, 4H). *NH groups were not detectable in this NMR experiment. LC-MS (ESI) m/z: [M+H]^+^ calcd. for C_20_H_27_F_3_N_3_O_3_S^+^, 446.17; found, 446.20.

*(1R,3S)-*3-((4-methyl-6-(methylamino)-1,3,5-triazine-2-yl)amino)-*N*-(4-(pent-4-yn-1-ylsulfon-amido)-2-(trifluoromethyl)benzyl)cyclohexane-1-carboxamide (**A2**): 2,4-dichloro-6-methyl-1,3,5-triazine (26 mg, 0.15 mmol, 1.0 eq) and methylamine (40% in water, 17 µL, 0.15 mmol, 1.0 eq) were combined in a 5 mL round bottom flask. A 1 M sodium hydroxide solution was added dropwise until a pH of 12 was reached. A solution of (*1R,3S*)-3-amino-*N*-(4-(pent-4-yn-1-ylsulfonamido)-2-(trifluoromethyl)benzyl)cyclohexane-1-carboxamide (68 mg, 0.15 mmol, 1.0 eq) in MeOH (1 mL) was then added and a pH of 10 was adjusted with a 1 M sodium hydroxide solution. The reaction mixture was stirred under reflux for 16 h. Afterwards, the solvent was removed under reduced pressure and the crude product was purified by flash chromatography (DCM:MeOH = 100:0 → 90:10). The product was obtained as a yellow solid in 20 % yield (15 mg, 0.0134 mmol). ^1^H NMR (400 MHz, DCM-d_2_): *δ* 7.56 (d, J = 2.0 Hz, 1H), 7.39 (d, J = 8.4 Hz, 1H), 7.33 -7.30(m, 1H), 4.58 -4.47 (m, 2H), 3.99 -3.97 (m, 1H), 3.27 -3.20 (m, 2H), 2.90 -2.86 (m, 2H), 2.33 (dt, J = 2.6 Hz, J = 6.8 Hz, 3H), 2.17 -3.13 (m, 2H), 2.04 -1.97 (m, 4H), 1.86 -1.80 (m, 2H), 1.40 -1.39 (m, 2H), 1.26 (s, 3H), 1.63 (d, J = 6.1 Hz, 1H), 0.89 -0.84 (m, 1H). *4 x NH were not detectable in this ^1^H NMR experiment. LC-MS (ESI) m/z: [M+H]^+^ calcd. for C_25_H_33_F_3_N_7_O_3_S^+^, 568.23; found, 568.35.

*N*-(pent-4-yn-1-yl)-4-(((1*r*,4*r*)-4-(3-(4-(trifluoromethoxy)-phenyl)ureido)cyclohexyl)oxy)-benzamide (**A3**): 4-[[*trans*-4-[[[[4-(trifluoromethoxy)phenyl]amino]carbonyl]amino]cyclohexyl]-oxy]benzoic acid (*t*-TUCB) (55 mg, 0.13 mmol, 1.0 eq), HATU (95 mg, 0.25 mmol, 2.0 eq) and TEA (0.18 mL, 1.25 mmol, 10.0 eq) were dissolved in DMF (1 mL) and stirred for 10 min. Subsequently, pent-4-yn-1-amine hydrochloride (17 mg, 0.14 mmol, 1.1 eq) was added and the reaction was stirred for 16 h at rt Afterwards, the mixture was diluted with EtOAc (10 mL) and washed with brine (2x 10 mL). The aqueous phase was extracted with EtOAc (3x 15 mL) and the combined organic phases were dried over MgSO_4_ and filtered. The solvent was removed under reduced pressure and the crude product was purified using semipreparative HPLC (method A). ^1^H NMR (300 MHz, DMSO-d_6_): δ 8.51 (s, 1H), 8.30 (t, J=5.5 Hz, 1H), 7.82 - 7.76 (m, 2H), 7.49 - 7.45 (m, 2H), 7.24 - 7.19 (m, 2H), 7.01 - 6.96 (m, 2H), 6.19 (d, J=7.4 Hz, 1H), 4.46 - 4.40 (m, 1H), 3.58 - 3.48 (m, 1H), 3.30 - 3.25 (m, 2H), 2.78 (t, J=2.6 Hz, 1H), 2.24 - 2.17 (m, 2H), 2.07 - 1.92 (m, 4H), 1.74 - 1.64 (m, 2H), 1.55 - 1.31 (m, 4H). ^19^F NMR (282 MHz, DMSO-d_6_): δ - 57.1. LC-MS (ESI) m/z: [M+H]^+^ calcd. for C_26_H_29_F_3_N_3_O_4_^+^, 504.21; found, 504.25.

*N*-(but-3-yn-1-yl)-4-(((1*r*,4*r*)-4-(3-(4-(trifluoromethoxy)phenyl)ureido)cyclohexyl)oxy)-benzamide (**A4**): 4- [[*trans*-4-[[[[4-(trifluoromethoxy)phenyl]amino]carbonyl]amino]cyclohexyl]-oxy]benzoic acid (*t*-TUCB) (20 mg, 0.045 mmol, 1.0 eq), HATU (35 mg, 0.09 mmol, 2.0 eq) and TEA (0.06 mL, 0.45 mmol, 10.0 eq) were dissolved in DMF (1 mL) and stirred for 10 min. Subsequently, but-3-yn-1-amine hydrochloride (5 mg, 0.05 mmol, 1.1 eq) was added and the reaction was stirred for 16 h at rt Afterwards, the mixture was diluted with EtOAc (10 mL) and washed with brine (2x 10 mL). The aqueous phase was extracted with EtOAc (3x 15 mL) and the combined organic phases were dried over MgSO_4_ and filtered. The solvent was removed under reduced pressure and the crude product was purified using semipreparative HPLC (method A). ^1^H NMR (300 MHz, DMSO-d_6_): δ 8.52 (s, 1H), 8.46 (t, J=5.5 Hz, 1H), 7.82 - 7.77 (m, 2H), 7.50 - 7.45 (m, 2H), 7.25 - 7.19 (m, 2H), 7.04 - 6.99 (m, 2H), 6.20 (d, J=7.6 Hz, 1H), 4.47 - 4.41 (m, 1H), 3.55 - 3.51 (m, 1H), 3.39 - 3.37 (m, 2H), 2.82 (t, J=2.5 Hz, 1H), 2.41 (dt, J=2.4, 7.1 Hz, 2H), 2.08 - 1.92 (m, 4H), 1.55 - 1.31 (m, 4H). ^19^F NMR (282 MHz, DMSO-d_6_): δ - 57.1. LC-MS (ESI) m/z: [M+H]^+^ calcd. for C_26_H_29_F_3_N_3_O_4_^+^, 409.19; found, 490.15.

*N*-(3-(1-(2-((2-(2,6-dioxopiperidin-3-yl)-3-oxoisoindolin-5-yl)oxy)ethyl)-1*H*-1,2,3-triazol-4-yl)propyl)-4-(((1*r*,4*r*)-4-(3-(4-(trifluoromethoxy)phenyl)ureido)cyclohexyl)oxy)benzamide (**P3**): In a microtube, sEH ligand **A3** (10.0 mg, 0.02 mmol, 1.0 eq) and 3-(6-(2-azidoethoxy)-1-oxoisoindolin-2-yl)piperidine-2,6-dione (6.5 mg, 0.02 mmol, 1.0 eq) were dissolved in DMSO (200 µL) and 25 µL of the respective aqueous solutions of CuSO_4_.5H_2_O (1.5 mg, 0.006 mmol, 0.3 eq) and sodium ascorbate (1.2 mg, 0.006 mmol, 0.3 eq) were added subsequently (solvent ratio DMSO: H_2_O = 4:1). The reaction mixture was stirred at rt for 16 h and then directly subjected to preparative HPLC purification (method A) to obtain **P3** as a white solid in 67% yield (11.0 mg, 13.2 µmol).^1^H NMR (400 MHz, DMSO-d_6_): δ 10.98 (s, 1H), 8.54 (s, 1H), 8.33 (t, *J* = 5.6 Hz, 1H), 7.98 (s, 1H), 7.81 (t, *J* = 11.8 Hz, 2H), 7.60 – 7.38 (m, 3H), 7.21 (ddd, *J* = 12.4, 10.8, 2.4 Hz, 4H), 6.99 (d, *J* = 8.9 Hz, 2H), 6.21 (d, *J* = 7.6 Hz, 1H), 5.10 (dd, *J* = 13.3, 5.1 Hz, 1H), 4.73 (t, *J* = 5.0 Hz, 2H), 4.48 (t, *J* = 5.0 Hz, 2H), 4.42 (dd, *J* = 8.8, 5.0 Hz, 1H), 4.30 (dd, *J* = 50.2, 17.0 Hz, 2H), 3.60 – 3.47 (m, 1H), 3.30 – 3.22 (m, 2H), 2.98 – 2.82 (m, 1H), 2.66 (t, *J* = 7.5 Hz, 2H), 2.59 (dd, *J* = 15.2, 2.1 Hz, 1H), 2.37 (qd, *J* = 13.2, 4.4 Hz, 1H), 2.09 – 1.90 (m, 5H), 1.88 – 1.79 (m, 2H), 1.55 – 1.31 (m, 4H). ^19^F NMR (282 MHz, DMSO-d_6_): δ - 57.1. ^13^C NMR (101 MHz, DMSO-d_6_): δ 173.3, 171.5, 168.4, 166.2, 160.1, 158.5, 154.9, 147.0, 142.4, 140.3, 135.1, 133.5, 129.5, 127.1, 125.1, 122.9, 122.1, 120.5, 119.5, 119.0, 115.3, 107.8, 74.6, 67.2, 52.2, 49.2, 47.6, 47.2, 31.7, 30.4, 30.1, 29.5, 23.2, 22.9. LC-MS (ESI) m/z: [M+H]^+^ calcd for C_41_H_44_F_3_N_8_O_8_^+^: 833.32; found: 833.15. HRMS (ESI): [M+H]^+^ calcd for C_41_H_44_F_3_N_8_O_8_^+^: 833.3228; found: 833.3248. Purity (HPLC-UV 254 nm): 99%. *t*_R_ (method A) = 8.8 min.

*N*-(3-(1-(2-(4-(2-(2,6-dioxopiperidin-3-yl)-1,3-dioxoisoindolin-5-yl)piperazin-1-yl)ethyl)-1*H*-1,2,3-triazol-4-yl)propyl)-4-(((1*r*,4*r*)-4-(3-(4-(trifluoromethoxy)phenyl)ureido)cyclohexyl)oxy)-benzamide (**P4**): In a microtube, sEH ligand **A3** (10.0 mg, 0.02 mmol, 1.0 eq) and 3-(6-(2-azidoethoxy)-1-oxoisoindolin-2-yl)piperidine-2,6-dione (6.5 mg, 0.02 mmol, 1.0 eq) were dissolved in DMSO (200 µL) and 25 µL of the respective aqueous solutions of CuSO_4_.5H_2_O (1.5 mg, 0.006 mmol, 0.3 eq) and sodium ascorbate (1.2 mg, 0.006 mmol, 0.3 eq) were added subsequently (solvent ratio DMSO: H_2_O = 4:1). The reaction mixture was stirred at rt for 16 h and then directly subjected to preparative HPLC purification (method B) to obtain **P4** as a yellow solid in 48% yield (8.7 mg, 9.51 µmol). ^1^H NMR (400 MHz, DMSO-d_6_): δ 11.08 (s, 1H), 8.72 (s, 1H), 8.32 (t, *J* = 5.6 Hz, 1H), 7.91 (s, 1H), 7.77 (d, *J* = 8.9 Hz, 2H), 7.66 (d, *J* = 8.5 Hz, 1H), 7.52 – 7.44 (m, 2H), 7.34 (d, *J* = 2.1 Hz, 1H), 7.27 – 7.19 (m, 3H), 6.98 (d, *J* = 8.9 Hz, 2H), 6.39 (d, *J* = 7.6 Hz, 1H), 5.07 (dd, *J* = 12.9, 5.4 Hz, 1H), 4.47 (t, *J* = 6.3 Hz, 2H), 4.44 – 4.37 (m, 1H), 3.60 – 3.48 (m, 1H), 3.41 (dd, *J* = 11.9, 6.9 Hz, 4H), 3.30 – 3.23 (m, 2H), 2.88 (ddd, *J* = 17.4, 14.2, 5.6 Hz, 1H), 2.79 (t, *J* = 6.4 Hz, 2H), 2.66 (dd, *J* = 9.7, 5.4 Hz, 2H), 2.60 (d, *J* = 2.7 Hz, 1H), 2.56 (dd, *J* = 8.9, 3.3 Hz, 5H), 2.09 – 1.98 (m, 3H), 1.92 (dd, *J* = 17.8, 8.6 Hz, 2H), 1.81 (ddd, *J* = 14.7, 11.1, 7.3 Hz, 2H), 1.42 (dq, *J* = 22.4, 9.9 Hz, 4H). ^19^F NMR (282 MHz, DMSO-d_6_) δ - 57.1. ^13^C NMR (101 MHz, DMSO-d_6_) δ 172.8, 170.1, 167.6, 167.5, 165.5, 163.4, 161.5, 159.7, 155.3, 154.5, 150.8, 146.2, 141.9, 139.9, 133.9, 128.9, 126.7, 124.9, 122.2, 121.6, 118.5, 118.3, 117.8, 114.8, 108.0, 74.3, 60.6, 56.9, 51.9, 48.9, 47.2, 47.2, 46.8, 31.0, 29.9, 29.7, 29.0, 22.7, 22.2. LC-MS (ESI) m/z: [M+H]^+^ calcd for C_45_H_50_F_3_N_10_O_8_^+^: 915.37; found: 915.55. HRMS (ESI): [M+H]^+^ calcd for C_45_H_50_F_3_N_10_O_8_^+^: 915.3759; found: 915.3773. Purity (HPLC-UV 254 nm): 99%. *t*_R_ (method A) = 6.3 min.

### Assay procedures

sEH HiBiT Lytic Assay: PROTAC-induced degradation of sEH was assessed using the previously established sEH-HiBiT lytic assay according to the published protocol.^14^ In brief, HeLa cells stably overexpressing the sEH-HiBiT fusion protein (herein referred to as HeLa^sEH-HiBiT^) were maintained in growth medium (DMEM (1X) medium with phenol red (Thermo Fisher Scientific, #41965-039) supplemented with 10% Corning® Fetal Bovine Serum (Corning®, 35-079-CV), penicillin (100 units/ml), streptomycin (100 μg/mL) (Gibco #15140), and 1 mM sodium pyruvate (Gibco #11360)) at 37°C and 5% CO2. In preparation for the assay, cells were harvested in growth medium, cell density was adjusted to 4·10^5^ cells/mL, and the cells were seeded into a 384-well TC plate (NuncTM white polystyrole, flat bottom, # 1262058) using a Multidrop combi (Thermo Fisher Scientific) at 50 µL/well resulting in 2000 cells per well. The plate was sealed with a semipermeable AeraSeal™ film (Sigma-Aldrich/Merck, #A9224) and cells were incubated for 24 h at 37 °C and 5% CO_2_. For the screening, crude PROTACs were tested with a final concentration of 300 nM in the assay. In a 96-Deepwell plate (nerbe plus, #04-072-0500), crude mixtures were diluted with DMSO to a concentration of 60 µM. Afterwards, the solutions were further diluted with growth medium to a concentration of 3.3 µM (5.5% DMSO). For dose-response measurements of the PROTACs **P3** and **P4**, dilution series were prepared in growth medium from DMSO stocks (5.5% DMSO). In the assay, 5 µL of the respective dilutions were added to the cells in triplicates for a final volume of 55 µL and a final DMSO concentration of 0.5%. The plate was centrifuged for 1 min at 300 rpm, then resealed with AeraSealM™ film, and the cells were incubated for 18 h at 37 °C and 5% CO_2_. For PROTACs **P3** and **P4**, the same procedure was additionally conducted with incubation times of 24 h, 3 h and 1 h. After incubation, cells were washed four times with DPBS using a HydrospeedTM plate washer device (Tecan) with a remaining volume of 10 µL in each well. Cell lysis was performed by adding 1 µL of Mammalian Lysis Buffer (Promega) to each well which was followed by centrifugation for 1 min at 300 rpm and incubation for 10 min at rt. Meanwhile, the Nano-Glo® substrate mix was freshly prepared from 1600 µL Nano-Glo® HiBiT Extracellular Buffer, 52 µL Nano-Glo® HiBiT Extracellular Substrate and 26 µL LgBiT Protein (all part of the Nano-Glo® HiBiT Extracellular Detection System Kit, Promega). Subsequently, 10 µL of the Nano-Glo® substrate mix were added to each well, followed by centrifugation for 1 min at 300 rpm. After incubation for 10 min at rt, the luminescence signal was detected using a Spark Multimode Microplate Reader (Tecan). The mean luminescence signal of each triplicate relative to the mean DMSO control signal was plotted against time or concentration in Prism 7.0 (GraphPad Software, Inc.). To determine the DC_50_ values of **P3** and **P4**, the normalized luminescence signals were plotted against the logarithmic compound concentration, and data analysis was performed using the nonlinear regression curve fit “log(Inhibitor) vs. Response –Variable slope (four parameters)” in Prism 7.0 and are calculated as mean ± SD.

For the viability assay, HeLa^sEH-HiBiT^ were maintained in growth medium (DMEM (1X) medium with phenol red (Thermo Fisher Scientific, #41965-039) supplemented with 10% Corning® Fetal Bovine Serum (Corning®, 35-079-CV), penicillin (100 units/ml), streptomycin (100 μg/mL) (Gibco #15140), and 1 mM sodium pyruvate (Gibco #11360)) at 37 °C and 5% CO_2_. The cell suspension was adjusted to a density of 200,000 cells/mL, and 30 μL/well, (equivalent to 6000 cells/well) were seeded into 96-well half area white PS flat bottom TC plates (Greiner bio-One, #675083). CuSO_4_^·^5 H_2_O and sodium ascorbate were prepared as dilution series in medium containing 2% DMSO. Each well received 5 μL of the respective dilution, resulting in the desired concentrations and a final DMSO concentration of 0.5%. As a performance control, 10 μM staurosporine was included in parallel. After 24 hours of treatment, 20 μL of CellTiter-Glo reagent (Promega #G7570) were added per well. The plate was protected from light and incubated for 30 minutes at rt before luminescence was measured using the standard attenuation protocol on a Spark Multimode Microplate Reader (Tecan). Mean of three technical replicates (N = 3) per concentration was normalized using wells with medium alone and wells with cells treated with only DMSO as 0 and 100% cell viability control. The livability rates are given as mean ± SD, calculated in Prism 7.0.

## Acknowledgements

This research was supported by Deutsche Forschungsgemeinschaft (Sachbeihilfe 530858826 to EP) and by the Proximity-inducing Drugs platform of the Fraunhofer Cluster of Immune Mediated Diseases. NL, SK and KH are grateful for support by the translational cancer program of the German Cancer Aid TACTIC (70115201). AK was supported by the Leistungszentrum Innovative Therapeutics (TheraNova) funded by the Fraunhofer Society and the Hessian Ministry of Science and Art.

## Contributions

Conceptualization: JS, EP, KH; Synthesis: JS, NL; biochemical assays: SB, JS, KH, LW; Writing the original draft: JS, KH; Editing: SK, AK; Manuscript review: all authors.

